# Cardiorespiratory mechanisms triggered during music perception in preterm infants and adults

**DOI:** 10.1101/2025.06.03.657691

**Authors:** L. Lavezzo, F. Barcos Munoz, D. Benis, J. Margaria, D. Grandjean, E.P. Scilingo, A. Vollenweider, S. Henriot, M. Nardelli, P. Hüppi, M. Filippa

**Affiliations:** Department of Information Engineering, University of Pisa; Division of Pediatric Intensive Care and Neonatology, Department of Women, Children and Adolescents, University Hospital of Geneva; Swiss Center for Affective Sciences, Department of Psychology and Educational Sciences, University of Geneva; Division of Development and Growth, Child and Adolescent Department, University of Geneva

**Author notes:** The co-last authors equally contributed to the manuscript (M. Nardelli, P. Hüppi, M. Filippa).

**Keywords:** Preterm newborns, music processing, heart rate variability, respiratory sinus arrhythmia

## Abstract

Preterm newborns’ autonomic response to dynamic auditory stimuli is poorly understood. To examine how cardiac and respiratory systems adjust their rhythms in response to music, we assessed 18 preterm infants (gestational age 37.73 ± 0.80 weeks) and 19 adults across music listening. Heart rate variability (HRV) dynamics and respiratory sinus arrhythmia (RSA), and cardiorespiratory coupling were analyzed. Both groups showed decreased high-frequency power in HRV during music listening. In preterm infants, RSA increased (p-value = 0.005), possibly suggesting a state of calm alertness, where the infant is physiologically prepared for interaction with environmental stimuli, while adults had decreased RSA (p-value = 0.003) and concomitant increased values of the low frequency/high frequency power ratio, possibly reflecting heightened alertness. While HRV responses to music were similar between preterms and adults, only the investigation of RSA and cardiorespiratory coupling measures revealed the delicate balance of autonomic dynamics in preterm newborns.

## INTRODUCTION

During hospitalization in the Neonatal Intensive Care Unit (NICU), preterm infants experience a phase of brain development in which activity dependent plasticity is prominent before reaching term-equivalent age (Kiss et al., 2014). During this period, they are exposed to many disturbing mechanical auditory stimuli, with significantly reduced exposure to human voices (Lahav & Skoe, 2014). This atypical auditory environment has increasingly been recognised to contribute to an increased risk of later adverse outcomes (Mayberry et al., 2002; Pineda et al., 2014).

Early enriching interventions are therefore investigated for supporting the development of preterm infants (Chung et al., 2020). Recent research indicates that environmentally mediated neuroprotective strategies can enhance their neurophysiological maturation, contributing to better neurobehavioral organization (Pickler et al., 2010). Among these, music-based interventions have gained increasing attention as a non-invasive means of fostering early structural and functional brain development (De Almeida et al., 2020; Lordier, Meskaldji, et al., 2019), promoting autonomic stability and enhancing overall well-being in preterm infants (Costa et al., 2022; Haslbeck et al., 2023; Mohan et al., 2021). In particular, recent studies also suggest that exposure to music and maternal singing can positively influence vital physiological functions, including heart rate, respiration, and stress regulation, by engaging the autonomic nervous system (Arnon et al., 2014; Barcos‐Munoz et al., 2024; Erdei et al., 2024; Filippa et al., 2024; Filippa et al., 2017).

Heart rate variability (HRV) has emerged as a key measure for assessing the impact of environmental stimuli such as music exposure on autonomic regulation in preterm infants (Foroushani et al., 2020). HRV reflects the dynamic balance between sympathetic and parasympathetic activity, providing valuable insights into the infant’s capacity for self-regulation, especially while attending to external stimuli.

Most studies investigating the effects of auditory stimulations on preterm infants’ cardiovascular system primarily focus on HRV itself, often using high-frequency (HF) power as a marker of parasympathetic activity (Yakobson et al., 2021). While HRV-HF power is commonly used, it does not directly quantify the respiration-related contributions to HRV. This is because it relies on spectral approximations rather than explicitly separating HRV into respiratory and non-respiratory components (Varon et al., 2019). Although traditional frequency-domain HRV parameters are essential, they do not clearly distinguish between the various mechanisms and pathways—both direct and indirect—that influence the heart (Varon et al., 2019). As a result, they limit our understanding of which cardiorespiratory mechanisms are triggered during f.e. music perception. This limitation is specifically relevant in preterm infants, where autonomic and respiratory control systems are in a delicate balance in the first weeks of life during the hospitalization period (Longin, Gerstner, Schaible, Lenz, & Koenig, 2006).

Indeed, to better understand the mechanisms underlying responses to music in the preterm population, it is important to investigate the interaction between two principal physiological systems—the cardiac and respiratory systems—under neural regulation, which is conventionally recognized through RSA (Angelone & Coulter JR, 1964; Song & Lehrer, 2003). Quantifying the respiration-driven modulations of the heart, particularly through RSA (Berntson et al., 1993), is thus crucial to gain new insights into music perception at the onset of the human auditory experience.

RSA is a key component of HRV that reflects the dynamic interplay between cardiac function and the respiratory cycle. While RSA is widely considered a non-invasive index of parasympathetic (vagal) cardiac control (Berntson et al., 1993), it is important to exercise caution when interpreting RSA as a direct or isolated measure of vagal tone, especially across different developmental stages. RSA magnitude can be influenced by multiple physiological and behavioral factors beyond vagal activity, including metabolic demands (e.g., Porges, 2007; Grossman & Taylor, 2007). In neonates and preterm infants, whose autonomic and respiratory systems are still maturing, these influences may vary considerably, potentially confounding straightforward interpretations of RSA as a pure marker of vagal function or interindividual differences. Consequently, although RSA provides valuable insight into autonomic reactivity and organization, our findings should be interpreted within the broader context of developmental variability and the multifactorial nature of cardiorespiratory coupling.

RSA manifests as an increase in heart rate during inspiration and a decrease during expiration, making it a well-established marker of vagal tone and autonomic regulation. A stronger vagal influence is linked to a more pronounced coupling between respiration and HRV, resulting in elevated RSA levels. In adults, RSA serves as the predominant mechanism of HRV modulation, where parasympathetic activity plays a crucial role, particularly in resting conditions (Hayano et al., 1996; Yasuma & Hayano, 2004). However, in preterm infants, the expression of RSA is less stable and more variable, reflecting the ongoing maturation of autonomic control mechanisms (Mulkey et al., 2021). This developmental immaturity suggests that their cardiac regulation is still evolving, making RSA a valuable measure for assessing the progressive refinement of vagal function during early life. Moreover, respiratory frequency may fall outside the high-frequency band (Varon et al., 2019), which is a concern for the application of standard frequency analysis on the signals acquired from newborn infants, whose respiratory rates are characterized by higher median values and greater variability (Fleming et al., 2011). To address these issues, research has applied specific methods that provide a direct and robust estimation of the respiratory-related HRV component based on orthogonal subspace decomposition (Varon et al., 2019), which has been identified as a reliable marker of RSA (Morales et al., 2021). This allows to distinguish between HRV changes due to respiration and those arising from other autonomic mechanisms.

In a variety of studies focusing on infants, RSA has been utilized as a physiological marker, owing to its non-invasive assessment capabilities through ECG and respiratory recordings (Abney et al., 2021; Dale et al., 2011; Feldman, 2006). However, research exploring its role in understanding the cardiorespiratory mechanisms underlying music perception across infancy remains limited.

One study addressing this question investigated how tonal and atonal music influence RSA in 40 mothers and their 3-month-old infants (Van Puyvelde et al., 2014). The tonal music fragment was based on harmonic structures resembling those found in mother-infant vocal interactions, whereas the atonal fragment lacked tonal organization. The findings revealed that infants exhibited significantly higher RSA levels when exposed to tonal music compared to both the atonal fragment and baseline conditions, which could be interpreted as enhanced vagal activity in response to tonal stimuli (Van Puyvelde et al., 2014). Interestingly, the study also found that RSA responses to tonal and atonal music differed between infants and mothers: while infants responded with increased RSA during the tonal segment, mothers showed distinct RSA patterns from their preterm infants. This divergence in response emerged when their cardiovascular systems did not exert mutual influence, highlighting the role of physiological synchronization in shaping their responses to musical stimuli.

In adults, research controlling for the respiratory component of RSA suggests that music’s effect on autonomic function is strongly influenced by its valence and arousing properties (Iwanaga & Tsukamoto, 1997). Most of the studies investigating physiological responses to music in adults are based or HR or HRV measures, but findings remain inconsistent, reflecting the complexity of autonomic regulation in reaction to emotional auditory stimuli (Kulinski et al., 2022). Arousing (Davis & Thaut, 1989) or calming effects (Mojtabavi et al., 2020) have been found, but the relationship between HR changes and musical characteristics is not always straightforward. Listening to stimulating music, as opposed to calming music, seems to be linked to a decrease in HRV, reflected in the reduction of both low-frequency and high-frequency HRV power, alongside an increase in HR (Koelsch & Jäncke, 2015). Additionally, when music elicits strong emotional responses heightened physiological arousal has been found (Rickard, 2004).

Building upon the role of different cardiorespiratory components, it is essential to acknowledge that additional mechanisms of cardiorespiratory phase coupling and synchronization represent different aspects of the cardiorespiratory interaction (Bartsch et al., 2012). Phase coupling refers to the phenomenon in which heart rate and respiration exhibit coordinated phase relationships, independent of RSA (Bartsch et al., 2014). Phase coupling is particularly relevant in preterm infants, whose developing cardiorespiratory systems exhibit distinctive patterns of coordination (Joshi et al., 2019; Lucchini et al., 2018). Studies suggest that the maturation of this coupling over time offers critical insights into the autonomic and cardiorespiratory system development (Solís-García et al., 2024). It is for this reason that we incorporated this additional measure in the present study to capture a more comprehensive understanding of the cardiorespiratory interaction.

Finally, while previous methodologies provide insights into the overall strength of heart-respiratory interactions, they do not account for directionality—i.e., whether the influence predominantly flows from respiration to the heart (classical RSA) or from the heart to respiration (non-respiratory HRV modulations). Given previous findings on developmental changes in autonomic control, investigating directionality in preterm newborns can better disentangle the differences between infants and adults across development (Joshi et al., 2019) (Mrowka et al., 2003). To address this, we applied Bivariate Phase-Rectified Signal Averaging (BPRSA) (Bauer et al., 2009), which allowed us to separately analyze cardiorespiratory coupling during heart rate acceleration and deceleration. Furthermore, we applied a novel approach, which for the first time studies the cross-correlation and mutual information of BPRSA series, to assess the main directionality between the dynamics of the cardiac and respiratory systems.

This study aims to explore the mechanisms underlying the cardiorespiratory response in preterm infants during music listening and to compare these dynamics with those of adults. To compare the two groups (18 preterm infants and 19 adults), we designed an experimental protocol consisting of three stages: two resting-state sessions without sound, each lasting 3 to 5 minutes—one at the beginning and one at the end—and a music elicitation phase in the middle, lasting 6 minutes. The composer, Andreas Vollenweider, created this sound environment for preemies, incorporating instruments such as the Indian flute (punji), harp, and bells, over a continuous background of harmonic sounds (Lordier, Loukas, et al., 2019). ECG signals were recorded from all participants throughout the experiment. By examining the dynamic interactions between traditional components of HRV and respiration across multiple levels, and by adding a specific directionality assessment, this investigation seeks to provide a comprehensive understanding of music-induced autonomic changes in preterm infants, identifying which specific mechanisms are modulated by the musical stimulus and how these changes are expressed in adults.

It is hypothesized that the high frequencies of the HRV in preterm infants and in adults will be affected during music listening, reflecting a shift in autonomic regulation. Furthermore, we predict that preterm infants will exhibit different patterns of cardiorespiratory coupling compared to adults in response to the music stimulus, which may be indicative of distinct stages in their cardiorespiratory and autonomic maturation. We also expect to find differences between preterms and adults in the main directionality of the coupling, shifting from heart-drive to respiration-driven with age as the RSA mechanism is fully developed in adulthood.

## RESULTS

### Music modulates preterm infant’s HRV

HRV analysis, the gold standard for autonomic assessment (“Heart rate variability,” 1996), has been applied to quantify autonomic activation during music stimulation. Specifically, we extracted frequency-domain indices to quantify the Autonomic Nervous System (ANS) activity: the high frequency (HF) power as index of purely parasympathetic activation, and the low-to-high frequency power ratio (LF/HF) as index of physiological arousal(“Heart rate variability,” 1996).

In preterm infants, a significant change in HF was observed throughout the experimental protocol (Friedman’s test, χ^2^(2) = 10.111, p = 0.006), with a notable decrease from the initial baseline to the music condition (V = 146, Bonferroni-corrected p-value = 0.020) and to the post condition (V = 152, Bonferroni-corrected p-value = 0.007). Figure 1 shows the boxplots of the HF power related to the three experimental conditions. No significant change was found for the LF/HF ratio (Friedman’s test, χ^2^(2) = 1.7778, p-value = 0.411). Additionally, to disentangle respiratory-related effects on the HRV dynamics from separate ANS control mechanisms, the Orthogonal Subspace Decomposition (OSP) approach was used ((Varon et al., 2019), see details in Section(STAR Methods). For each frequency band (HF and LF), we extracted two components from HRV power spectrum: a respiratory component related to respiration (hereinafter RespComp), and a residual component (hereinafter Residual) associated to sympathetic and vagal modulations unrelated to breathing functions.

**Figure 1.**
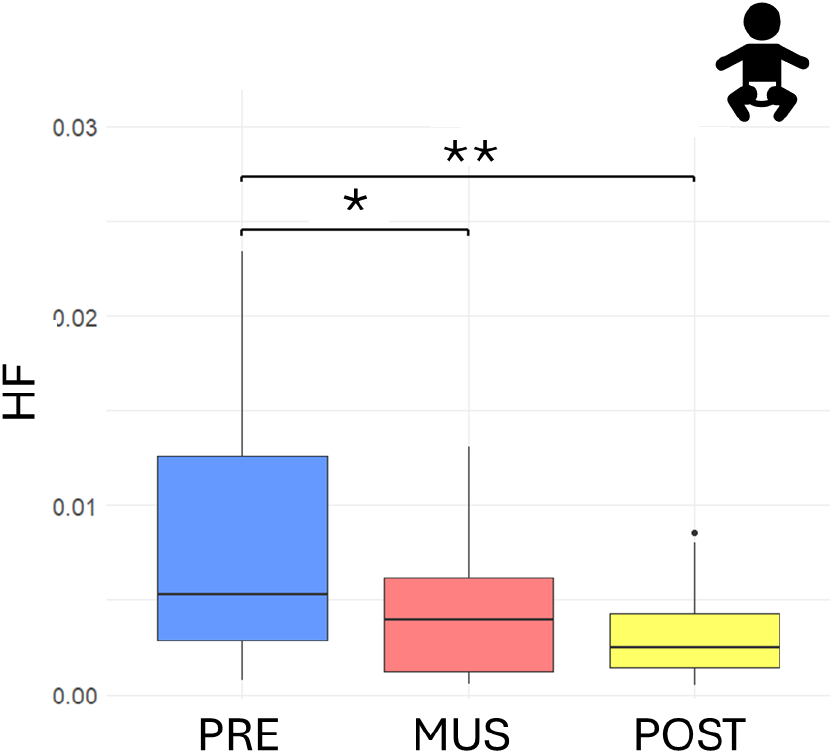
Boxplot representing the HF power for preterm infants (n = 18) before (blue) during (red) and after (yellow) music stimulation. Significance thresholds are represented by “* = p <0.05”, “** = p < 0.01”.

Concerning the analysis of the two respiratory-related components of HRV frequency domain features, the HF RespComp showed an increasing trend during the protocol (Friedman’s χ^2^(2) = 7, p-value=0.030). However, this change was not statistically significant after correcting the p-values for multiple comparisons (pre vs. music V = 44, adjusted-p = 0.221; pre vs. post V = 52, adjusted p = 0.462; mus vs. post V = 84; adjusted-p = 1). No significant change has been reported for the LF/HF RespComp.

The results obtained for the HF Residual were consistent with the standard analysis, indicating a significant decreasing trend for HF (Friedman test, χ^2^(2) = 12.333, p-value = 0.002). Subsequent pairwise comparisons revealed a significant difference between the initial baseline and the music condition (V = 147, Bonferroni-corrected p-value = 0.017), and between the initial baseline and the post condition (V = 152, Bonferroni-corrected p-value = 0.007). No significant difference was reported between the music condition and the post condition. The trends of the values of HF Residual in the three sessions are presented in Figure 2.

**Figure 2.**
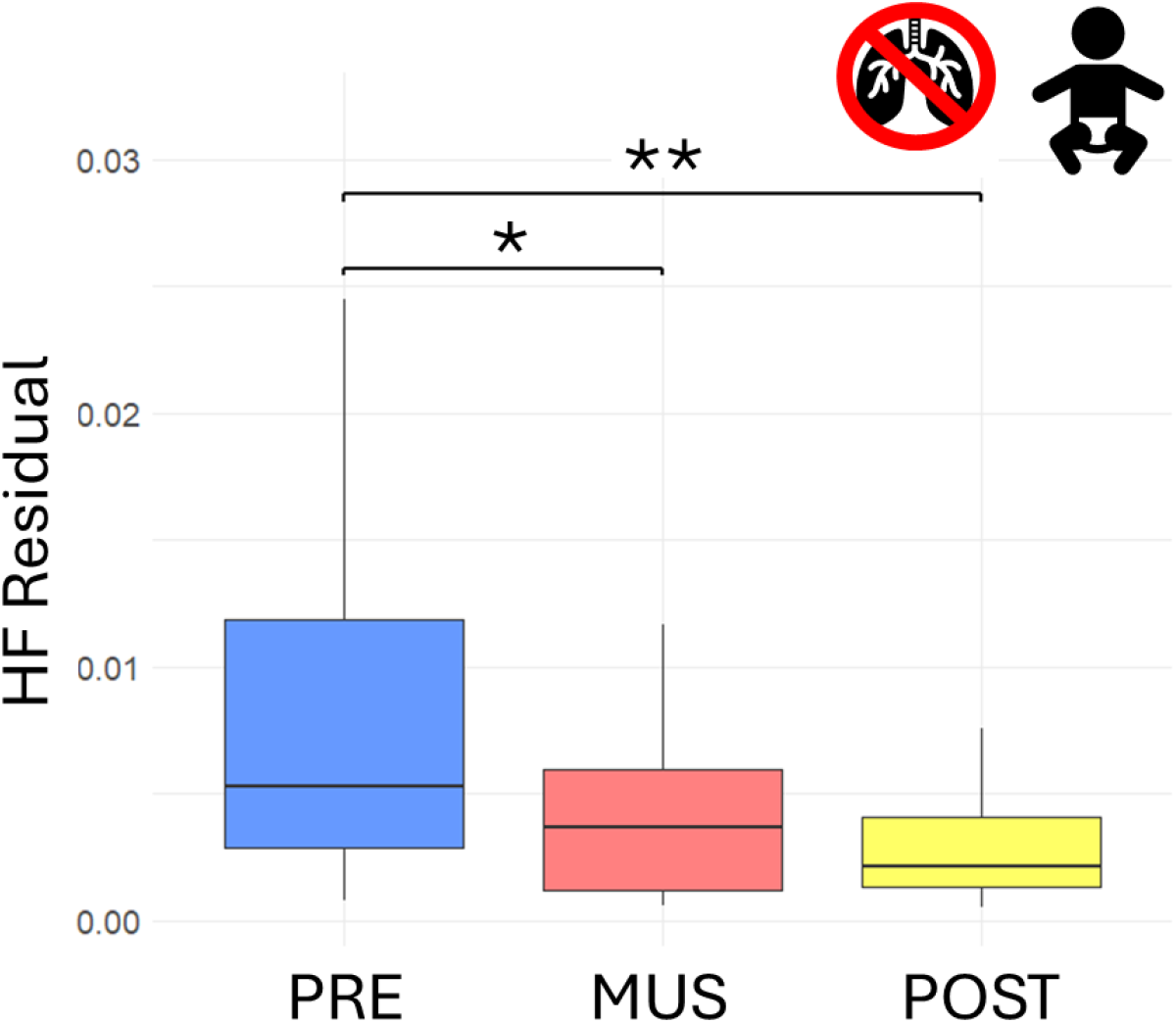
Boxplot representing the HF Residual component for preterm infants (n = 18) before (blue) during (red) and after (yellow) musical stimulation. Significance thresholds are represented by “* = p <0.05”, “** = p < 0.01”.

**Table 1.** Summary of metrics values obtained for infants during before and after the music protocol.

Infants
| Metric | Description | pre |  | music |  | post |  |
| --- | --- | --- | --- | --- | --- | --- | --- |
|  |  | mean | std | mean | std | mean | std |
| **HF** | Power in the high frequency band [0.2-1.5] | 0,008204 | 0,007212 | 0,004539 | 0,003851 | 0,00321 | 0,002383 |
| **LF/HF** | Ratio of low-to-high frequency power | 9,779873 | 5,448994 | 11,04703 | 5,370513 | 9,37231 | 7,833609 |
| **HF RespComp** | Power in the high frequency band [0.2-1.5] for respiratory component | 0,00023 | 0,000613 | 0,000352 | 0,000725 | 0,00051 | 0,001112 |
| **LF/HF RespComp** | Ratio of low-to-high frequency power for respiratory component | 0,329849 | 1,273773 | 0,119846 | 0,216701 | 1,405746 | 3,744098 |
| **HF Residual** | Power in the high frequency band [0.2-1.5] for residual component | 0,008136 | 0,007163 | 0,004194 | 0,003463 | 0,002812 | 0,001931 |
| **LF/HF Residual** | Ratio of low-to-high frequency power for residual component | 9,541607 | 5,277584 | 11,3398 | 5,356435 | 9,448729 | 7,162101 |

### Music modulate adult’s HRV

In the group of adults, a statistically significant change in HF power was observed (Friedman’s test, χ^2^(2) = 9.7895, p-value = 0.007), with a marked decrease from the initial baseline to the music condition (V = 160, Bonferroni-corrected p-value = 0.021). Additionally, the LF/HF ratio values exhibited significant changes across the three phases (χ^2^(2) = 15.579, Friedman’s test, p-value = 4.141e-4), showing a significant increase from the initial baseline to the music condition (V = 25, Bonferroni-corrected p-value = 0.010) and from the initial baseline to the post condition (V = 23, Bonferroni-corrected p-value = 0.007). The results are displayed in Figure 3.

**Figure 3.**
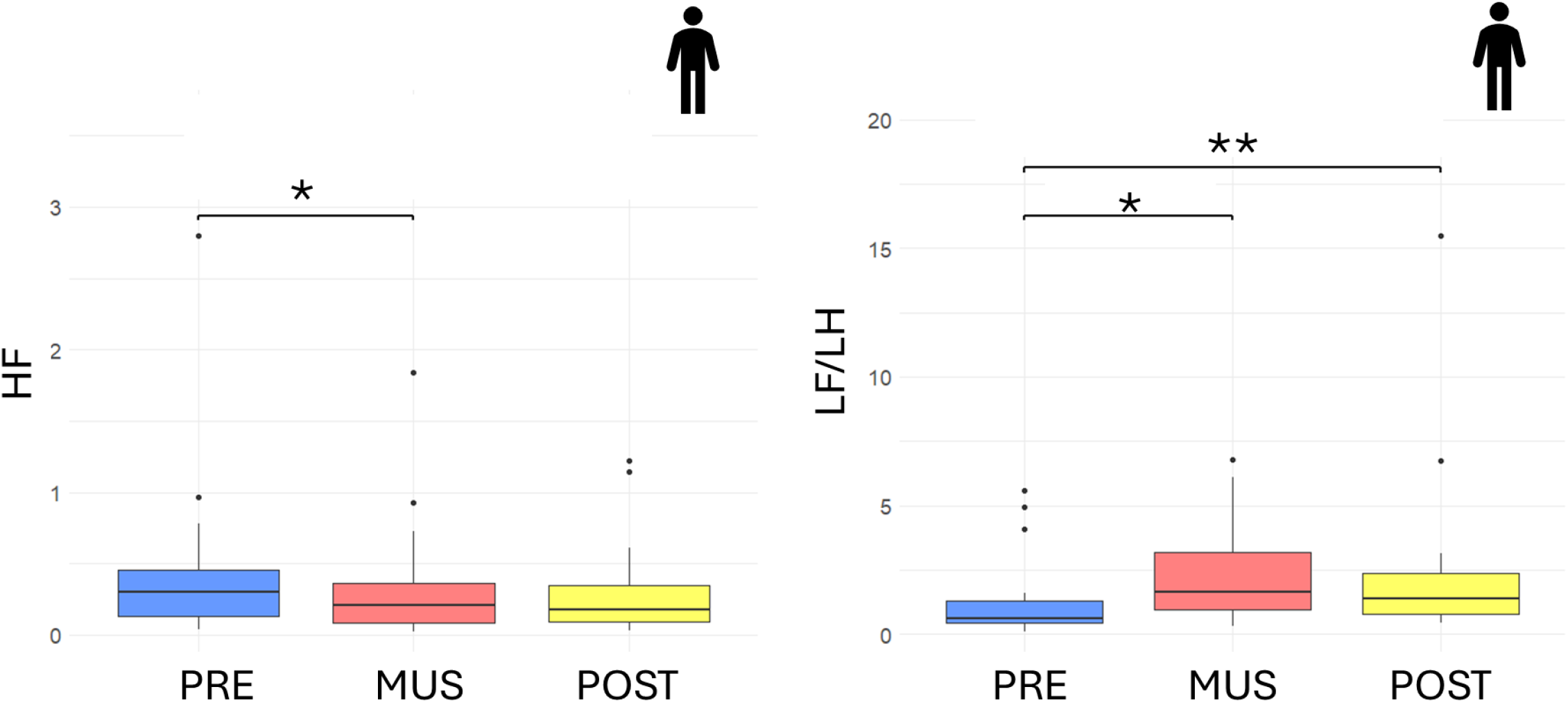
Boxplot representing the HF power (left) and the HF/LF ratio (right) for adults (n = 19) before (blue) during (red) and after (yellow) musical stimulation. Significance thresholds are represented by “* = p <0.05”, “** = p < 0.01”.

The respiratory-related components exhibited significant variations, particularly in HF RespComp (Friedman’s test, χ^2^(2) = 14, p-value = 0.009e-1). Post-hoc analyses revealed that HF RespComp was significantly higher in the initial baseline compared to the music condition (V = 181, Bonferroni-corrected p-value = 0.380e-3), and to the post condition (V = 169, Bonferroni-corrected p = 0.508e-2), indicating a significant reduction in respiratory-driven vagal modulation during music exposure. The boxplots reporting the median values of HF RespComp are presented in Figure 4. No significant differences were observed in HF Residual (Friedman test, χ^2^(2) = 0.73684, p-value = 0.691) in the three conditions.

**Figure 4.**
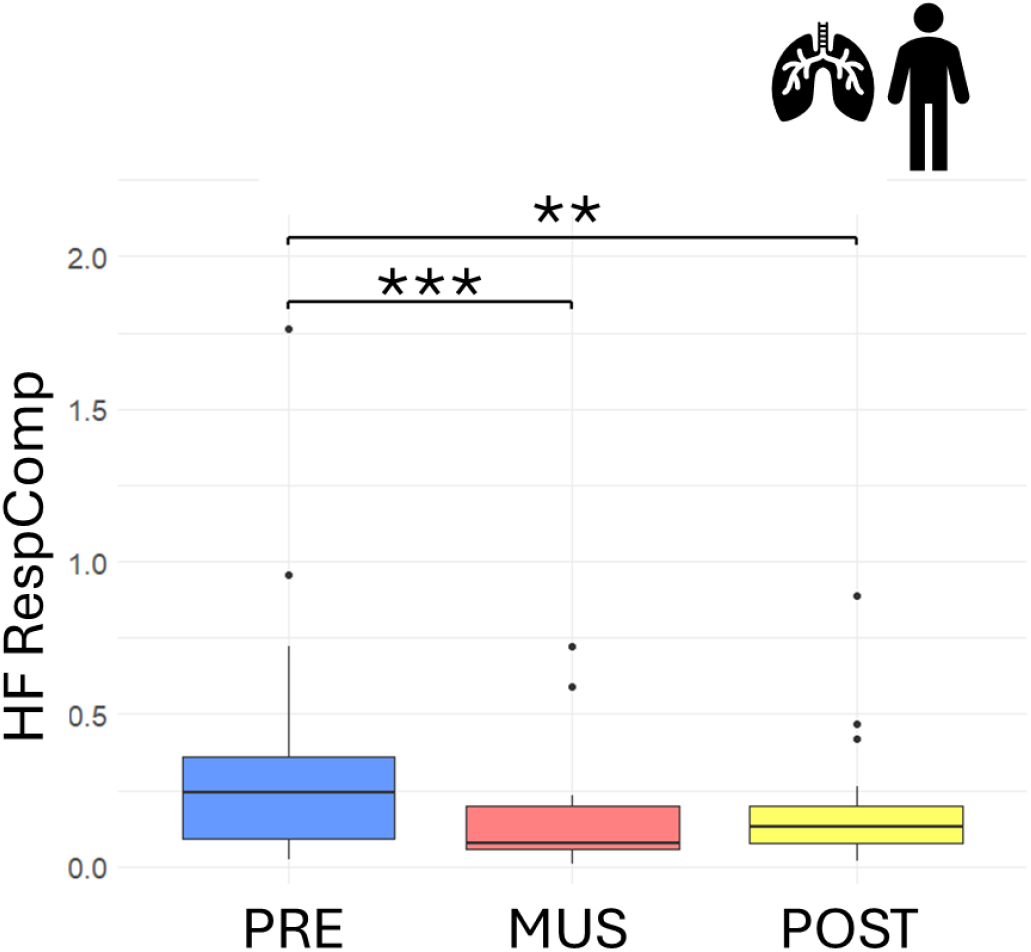
Boxplot representing the HF power of the respiratory component for adults (n = 19) before (blue) during (red) and after (yellow) musical stimulation. Significance thresholds are represented by “* = p <0.05”, “** = p < 0.01”.

**Table 2.**
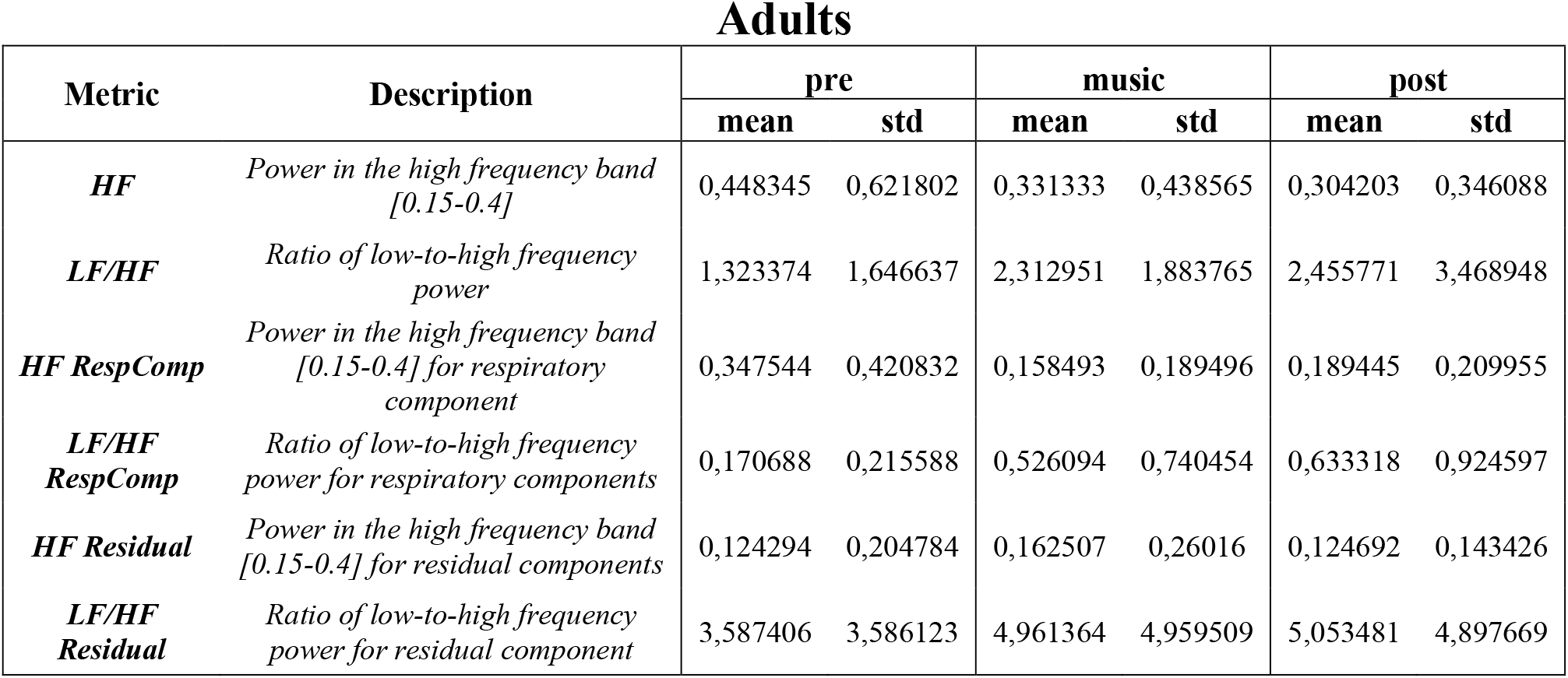
Summary of metrics values obtained for adults during before and after the music protocol.

### Music increases RSA and cardiorespiratory coupling in hospitalized preterm infants

Cardiorespiratory coupling has been assessed in its different aspects specifically, to effectively unveil the aspects of cardiorespiratory interactions in preterm infants and adults, we applied three main methodologies: the Orthogonal Subspace Projection (OSP) (Varon et al., 2019), the Mean Phase Coherence (MPC) (Schäfer et al., 1998), and the Bivariate Phase-Rectified Averaging (BPRSA) (Bauer et al., 2009).

Applying the OSP approach, we computed P_x_, i.e., the percentage of the HRV power related to the respiration component with respect to the total power, to determine the extent to which changes in HRV can be attributed to respiration (Varon et al., 2019), thereby serving as a more precise index of RSA (Morales et al., 2021).

The MPC directly quantifies the synchronicity between the phases of the two time-series regardless of the amplitude (Lanata et al., 2016).

In the infant population, P_x_ demonstrated a significant increasing trend across the different phases (Friedman’s test, χ^2^(2) = 10.778, p-value = 0.005). In particular, the strength of cardiorespiratory coupling increased significantly from the initial baseline to the post condition (V = 18, Bonferroni-corrected p-value = 0.006). The MPC displayed a similar increasing trend (Friedman’s test, χ^2^(2) = 6.3333, p-value = 0.042), however multiple comparisons were not significant after Bonferroni’s correction between the initial and the post condition (V = 37, Bonferroni-corrected p-value = 0.100).

The boxplots of P_x_ distributions in the three experimental sessions in preterm infants are displayed in Figure 5.

**Figure 5.**
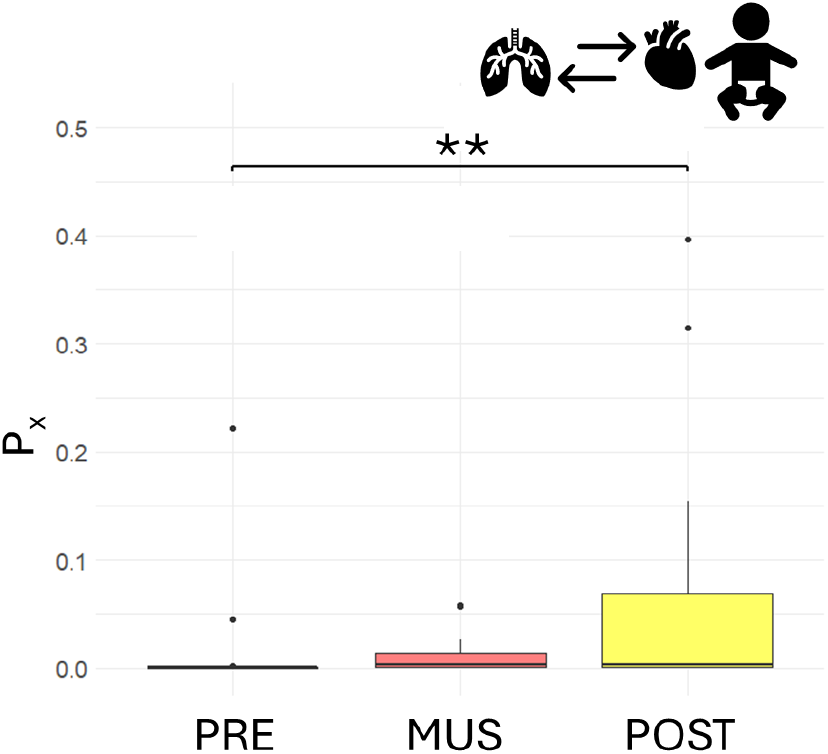
Boxplot representing the P_x_, for infants before (blue) during (red) and after (yellow) musical stimulation. Significance thresholds are represented by “* = p <0.05”, “** = p < 0.01”.

The BPRSA technique quantifies quasi-periodic oscillations in a time series due to interactions with another series. Coupling between signals has been measured using a linear metric, the maximum value of the cross-correlation function (max_xcorr), and a nonlinear marker, the Mutual Information (MI).

The BPRSA was used to firstly quantify the strength of the influence of RR series changes on the respiration using heart rate acceleration as anchor points. Relevant trends are represented in Figure 6. The MI values displayed an increasing trend during the experiment (Friedman’s test, χ^2^(2) = 11.444, p-value =0.003), with the multiple comparison tests being statistically significant between the initial and the post condition (V = 10, Bonferroni-corrected p-value=0.980e-3). The max_xcorr metrics computed between the two series reported a matching increasing trend however the difference is not significant (Friedman’s test, χ^2^(2) = 5.3333, p-value =0.069).

**Figure 6.**
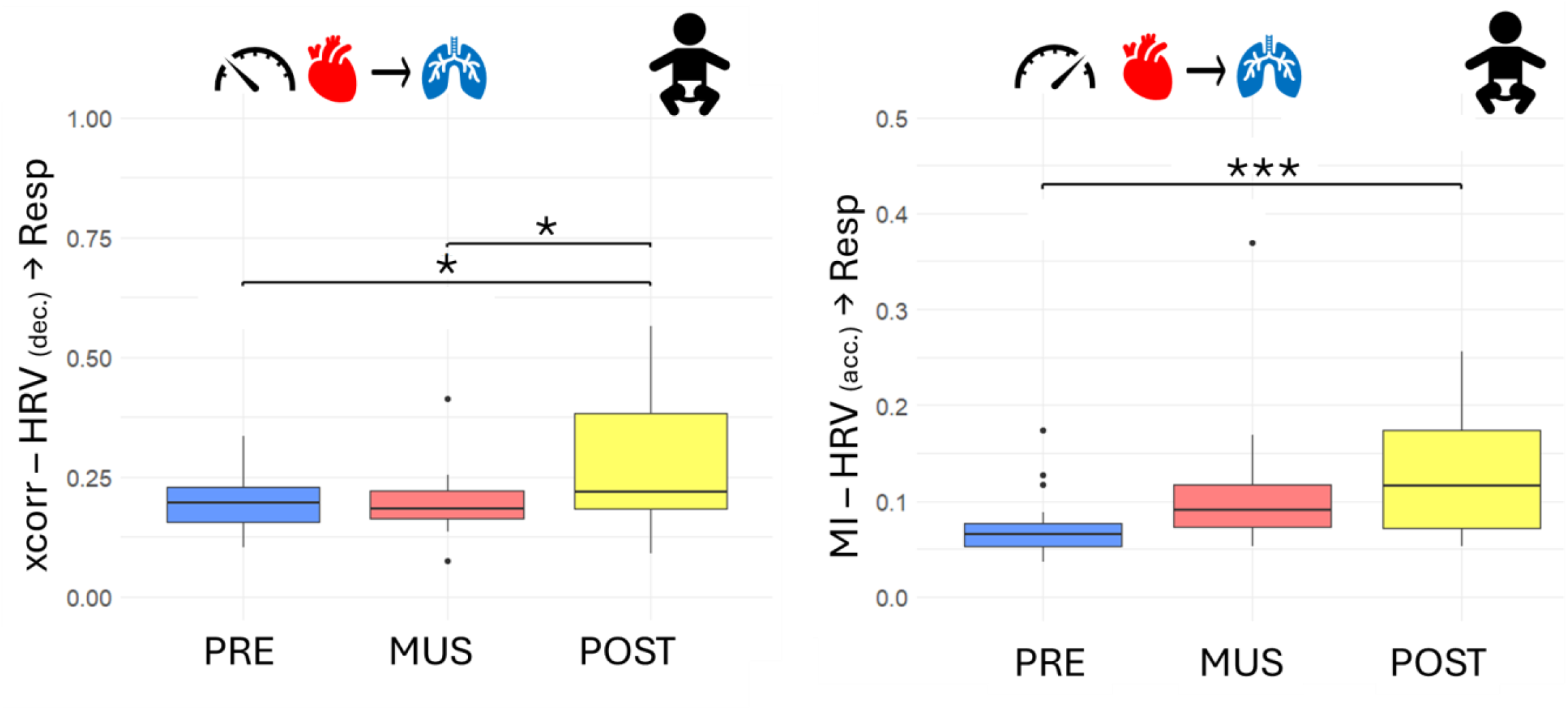
Boxplot depicting the BPRSA results for infants in NICU before (blue) during (red) and after (yellow) musical stimulation. On the right the relationship between the heart acceleration and the respiration, on the left the relationship between the heart deceleration and the respiration. Significance thresholds are represented by “* = p <0.05”, “** = p < 0.01”.

Concerning the effects investigated during the deceleration of the heart rate, a significant difference (Friedman’s test, χ^2^(2) = 7, p-value =0.030) was found in the post condition, which displayed values significantly higher than both the initial baseline (V = 30, Bonferroni-corrected p-value = 0.042) and the music condition (V = 27, Bonferroni-corrected p-value = 0.027). No significant differences are reported for the MI values after the multiple comparisons’ correction (Friedman’s test, χ^2^(2) = 6.3333, p-value =0.042).

BPRSA coupling in the direction going from the respiration to the heart during minimum and maximum impedance presented no statistically significant change during our experimental paradigm.

**Table 3.** Summary of cardiorespiratory coupling assessment for infants during before and after the music protocol.

Infants
| Metric | Description | pre |  | music |  | post |  |
| --- | --- | --- | --- | --- | --- | --- | --- |
|  |  | mean | std | mean | std | mean | std |
| **$P_x$** | *the power related to the respiration component with respect to the total power of the HRV signal* | 0,015174 | 0,052612 | 0,011649 | 0,01833 | 0,062765 | 0,115133 |
| **MPC** | *Mean Phase coherence between HRV & Respiration* | 0,051738 | 0,039092 | 0,07347 | 0,066613 | 0,095348 | 0,085693 |
| **MI - HRV (acceleration) -> Resp** | *Mutual information between trigger (HRV acceleration) & target (respiration)* | 0,073218 | 0,034821 | 0,109471 | 0,071592 | 0,127381 | 0,062722 |
| **MI - HRV (deceleration) -> Resp** | *Mutual information between trigger (HRV deceleration) & target (respiration)* | 0,079148 | 0,032712 | 0,113008 | 0,060013 | 0,102515 | 0,058334 |
| **xcorr - HRV (acceleration) -> Resp** | *Cross-correlation between trigger (HRV acceleration) & target (respiration)* | 0,206282 | 0,081108 | 0,21941 | 0,081268 | 0,293482 | 0,182323 |
| **xcorr - HRV (deceleration) -> Resp** | *Cross-correlation between trigger (HRV deceleration) & target (respiration)* | 0,198205 | 0,066926 | 0,197108 | 0,068263 | 0,290639 | 0,14881 |
| **MI - Resp (max) -> HRV** | *Mutual information between trigger (Respiration max) & target (HRV)* | 0,087447 | 0,039752 | 0,139976 | 0,106918 | 0,120287 | 0,099248 |
| **MI - Resp (min) -> HRV** | *Mutual information between trigger (Respiration min) & target (HRV)* | 0,089159 | 0,028408 | 0,13978 | 0,115719 | 0,125236 | 0,100091 |
| **xcorr - Resp (max) -> HRV** | *Cross-correlation between trigger (Respiration max) & target (HRV)* | 0,121213 | 0,078226 | 0,186635 | 0,125617 | 0,205467 | 0,185825 |
| **xcorr - Resp (min) -> HRV** | *Cross-correlation between trigger (Respiration min) & target (HRV)* | 0,134169 | 0,107247 | 0,197699 | 0,13704 | 0,195232 | 0,181676 |

### Music decreases RSA and cardiorespiratory coupling in adults

In the adult population, P_x_ showed a significant decrease (Friedmans’ test χ^2^(2) = 11.789, p-value = 0.003) from initial baseline to the music condition (V = 173, Bonferroni’s corrected p-value pre vs. music = 0.002) and between initial baseline and post condition (V = 164, Bonferroni’s corrected p-value pre vs. music = 0.011). No significant differences were reported between the music condition and the post condition. The related boxplots are displayed in Figure 7. The MPC results are consistent, showing the same marked decrease (Friedmans’ test, χ^2^(2) =, 15.895 p-value = 0.353 e-3), from the initial resting state session to the music condition (V = 179, Bonferroni’s corrected p-value pre vs. music = 0.630e-3) and between the initial baseline and post condition (V = 175, Bonferroni’s corrected p-value = 0.157 e-2).

**Figure 7.**
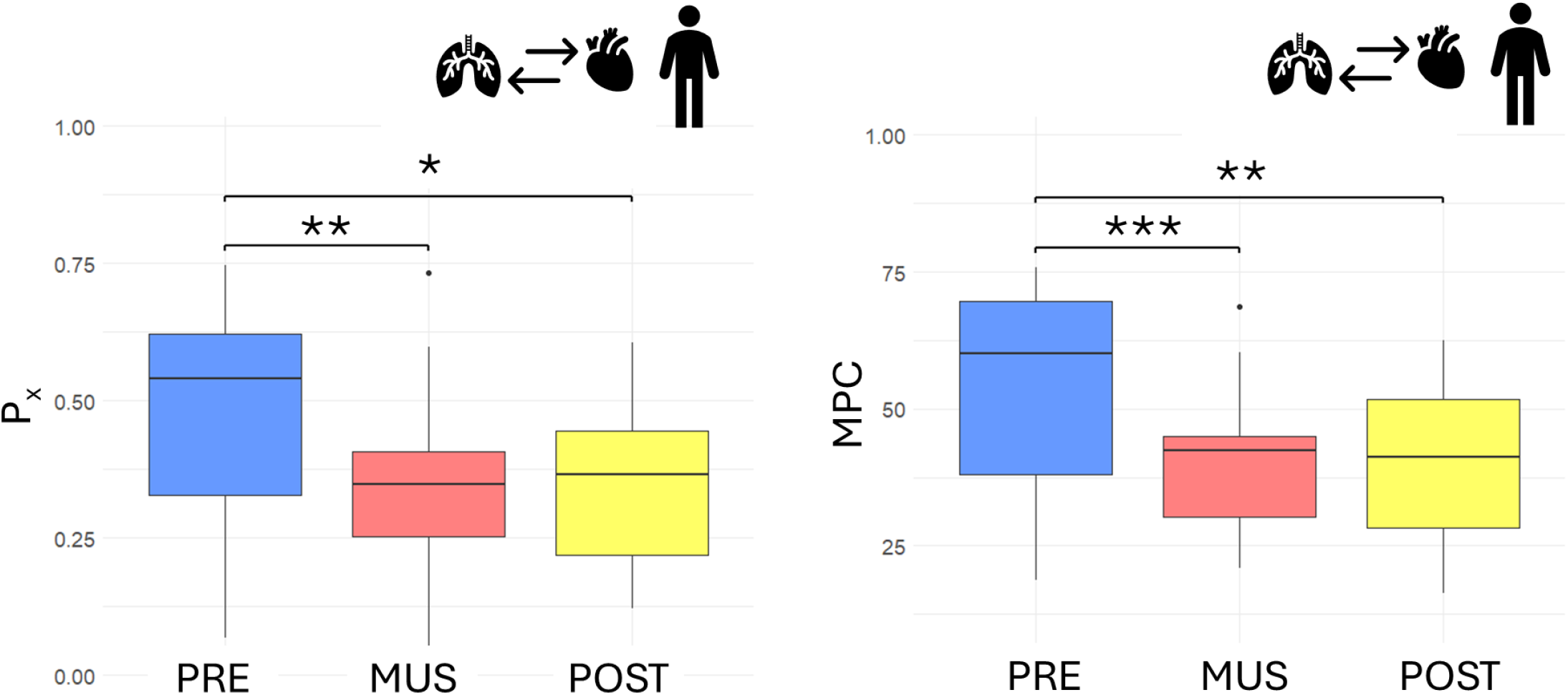
Boxplot depicting the P_x_ (left)and MPC (right) results for adults before (blue) during (red) and after (yellow) musical stimulation. Significance thresholds are represented by “* = p <0.05”, “** = p < 0.01”.

Regarding BPRSA, no significant change has been detected for any parameters (Friedmans’ test, max in cross correlation RR to RESP during heart rate acceleration χ^2^(2) = 3.2632, p-value = 0.195; during heart rate deceleration χ^2^(2) = 4.5263, p-value = 0.104; MI during heart rate acceleration χ^2^(2) = 2.2105, p-value = 0.331; during heart rate deceleration χ^2^(2) = 0.42105, p-value = 0.810). The trends of max_xcorr are shown to be consistent with previously reported results.

**Table 3.** Summary of cardiorespiratory coupling assessment for adults during before and after the music protocol.

| Metric | Description | pre |  | music |  | post |  |
| --- | --- | --- | --- | --- | --- | --- | --- |
|  |  | mean | std | mean | std | mean | std |
| $P_x$ | *the power related to the respiration component with respect to the total power of the HRV signal* | 0,480474 | 0,187891 | 0,338614 | 0,161297 | 0,351352 | 0,139318 |
| $MPC$ | *Mean Phase coherence between HRV & Respiration* | 0,554645 | 0,174877 | 0,399883 | 0,148792 | 0,392966 | 0,136645 |
| **MI - HRV (acceleration) -> Resp** | *Mutual information between trigger (HRV acceleration) & target (respiration)* | 0,336608 | 0,367421 | 0,232035 | 0,189881 | 0,253661 | 0,22115 |
| **MI - HRV (deceleration) -> Resp** | *Mutual information between trigger (HRV deceleration) & target (respiration)* | 0,326791 | 0,320487 | 0,247504 | 0,22466 | 0,255173 | 0,251278 |
| **xcorr - HRV (acceleration) -> Resp** | *Cross-correlation between trigger (HRV acceleration) & target (respiration)* | 0,788823 | 0,203796 | 0,695743 | 0,17845 | 0,706892 | 0,209103 |
| **xcorr - HRV (deceleration) -> Resp** | *Cross-correlation between trigger (HRV deceleration) & target (respiration)* | 0,799131 | 0,193346 | 0,710285 | 0,181315 | 0,682189 | 0,218088 |
| **MI - Resp (max) -> HRV** | *Mutual information between trigger (Respiration max) & target (HRV)* | 0,316509 | 0,32179 | 0,317323 | 0,259399 | 0,255873 | 0,256091 |
| **MI - Resp (min) -> HRV** | *Mutual information between trigger (Respiration min) & target (HRV)* | 0,364372 | 0,359241 | 0,319997 | 0,268196 | 0,276864 | 0,323218 |
| **xcorr - Resp (max) -> HRV** | *Cross-correlation between trigger (Respiration max) & target (HRV)* | 0,845897 | 0,13509 | 0,832187 | 0,14833 | 0,809727 | 0,166206 |
| **xcorr - Resp (min) -> HRV** | *Cross-correlation between trigger (Respiration min) & target (HRV)* | 0,873634 | 0,083933 | 0,828669 | 0,159192 | 0,795186 | 0,181538 |

### Main directionality of influence between cardiac and respiratory dynamics

To evaluate differences in the directionality of interactions, we used a novel approach based on the computation of cross-correlation and mutual information function on the BPRSA trigger and the target series (see Methods). The BPRSA indices of coupling strength from HRV to respiration during heart rate acceleration and deceleration were compared with those from respiration to HRV during minimum and maximum impedance, for each phase.

For premature infants, the analysis compounded of cross-correlation values assessing the strength of interactions between HRV and respiration across conditions, comparing the two possible directions of interaction, revealed significant differences in coupling direction. A stronger coupling from HRV to respiration was observed during the initial baseline (V = 594, Bonferroni-corrected p-value = 2.978 e^−5^) and the post condition (V = 532, Bonferroni-corrected p-value = 0.004). However, during music stimulation, no significant differences were found (V = 378, Bonferoni-corrected p-value = 1). Trends and relevant directions are depicted in Figure 8.

**Figure 8.**
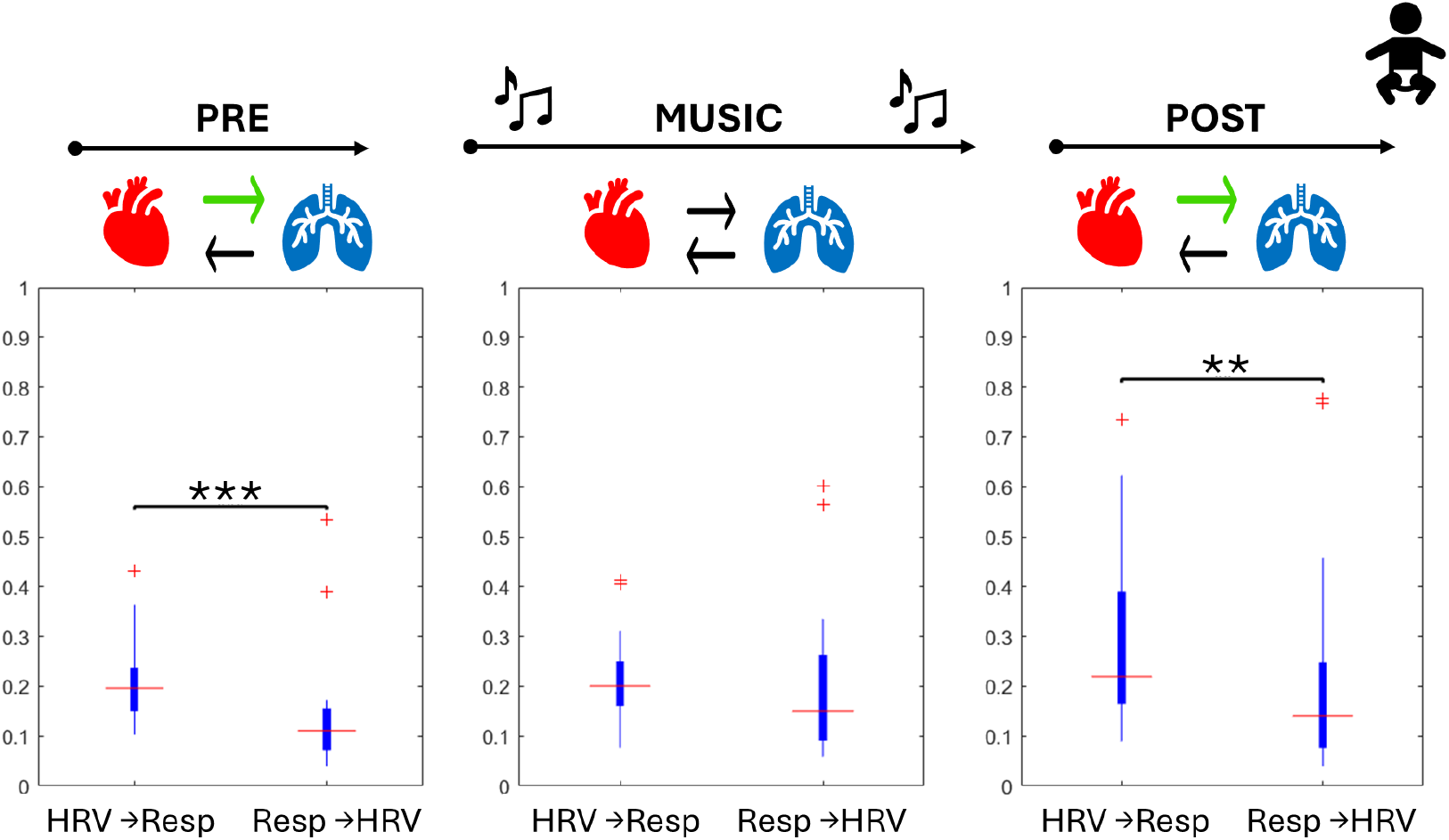
Directionality analysis for infants in NICU during the protocol. Boxplots depict the aggregation of cross correlation BPRSA values for each direction. Significance thresholds are represented by “* = p <0.05”, “** = p < 0.01”. In the top line protocol phases are represented, and arrow depicting a significantly stronger directionality is highlighted in yellow.

In adults, the analysis of cross-correlation values revealed significant differences in the strength of coupling directions (HRV to respiration vs. respiration to HRV) across all phases of the protocol, with stronger coupling from respiratory activity to heart activity. This difference was observed during the initial baseline (V = 189, Bonferroni-corrected p-value = 0.0228), the music phase (V = 84, Bonferroni-corrected p-value = 2.376 e^−5^), and the post condition (V = 131, Bonferroni-corrected p-value = 0.897 e^−3^).

For both preterm infants and adults, the mutual information-based approach did not highlight statistically significant differences between the various phases of the experimental protocol, expect for the music elicitation phase in adults were a consistent respiration to HRV flow was found.

## DISCUSSION

The goal of our study was to explore how the cardiac and respiratory systems adjust their rhythms and dynamics to a music stimulus in preterm newborns compared to adults. In particular, we aimed to examine different aspects of cardiovascular coupling to disentangle the underlying physiological processes that change during development.

To this end, we analysed the cardiac and cardiorespiratory responses of preterm infants and adults before, during, and after listening to a 6-minute music composed for preemies by Andreas Vollenweider (Lordier, Loukas, et al., 2019). HRV dynamics and RSA were assessed using frequency-domain analysis, cardiorespiratory phase coupling was measured by using the MPC method. Strength in the directionality was computed using the BPRSA approach.

The findings suggest that music listening has distinct effects on HRV in premature infants and adults. In preterm infants, music decreases the HF component, with the change attributed to the residual component, which is not linearly related to respiration. In contrast, music in adults decreases HF and increases the LF/HF ratio, with these changes being linearly related to respiration. Additionally, the primary directionality of interaction for preterm infants is from heart to respiratory dynamics, except during music stimulation, whereas for adults, the main directionality is from respiration to heart rhythm, consistently across all phases.

Firstly, the observed decrease in the high frequencies of the HRV alongside the increase in RSA suggests a shift towards emotional engagement and attentive state during music listening.

Across development, RSA is associated with vagal tone, which plays a crucial role in regulating attentional focus, emotional engagement, and social interaction (Porges & Carter, 2010). During dyadic interactions, an increase in RSA may indicate a state of calm alertness, where the infant is physiologically prepared for interaction with the environment (Feldman, 2012), including auditory stimuli such as music. Preterm infants often have immature autonomic regulation, which affects their ability to modulate arousal and attention. A shift toward greater RSA suggests that music or other external stimuli could be enhancing their self-regulatory capacity, promoting engagement with sensory input. Thus, in preterm newborns, a decrease in HRV HF power with a concurrent increase in RSA may suggest a reorganization of autonomic function toward a more focused or engaged state, rather than passive relaxation (Bazhenova et al., 2001). This could indicate a developmental adaptation where music or other stimuli help regulate their autonomic functions, potentially supporting early social and cognitive development.

Although RSA is predominantly modulated by parasympathetic activity, and increases in RSA are generally interpreted as reflecting enhanced vagal function, it is important to acknowledge that RSA is also influenced by respiratory patterns, metabolic state, and developmental factors. Nevertheless, empirical evidence supports the interpretation of higher RSA as an indicator of improved autonomic regulation and physiological stability. Groome et al. (1999) demonstrated that greater vagal tone is associated with more efficient homeostatic regulation in fetuses, while longitudinal human studies have shown that early improvements in RSA, particularly during sensitive periods and in relation to maternal contact (e.g., kangaroo care), predict more favourable developmental trajectories—including better maternal attachment behaviour, enhanced mother– child reciprocity at 10 years, improved executive functioning, and more adaptive autonomic profiles (Feldman et al., 2014).

Indeed, from an integrated psychophysiological point of view, RSA is one of the responses that are related to arousal-induced responses in social situations (Friedman, 2007). It has been speculated that during low demands for social engagement, vagal tone increases and HR decreases, allowing the body to focus on internal processes. By contrast, during challenging situations, vagal tone decreases and HR increases to prepare for environmental participation and self-regulation (Porges et al., 1994). However recent theoretical perspectives (Grossman, 2024; Ritz, 2024) emphasize that RSA is an indirect and imperfect proxy for parasympathetic influence, particularly on heart rate, and is closely tied to ventilatory-circulatory coordination. As such, interpreting RSA data necessitates careful consideration of respiratory parameters and acknowledgment that RSA does not exclusively reflect overall vagal tone or specific psychological constructs (Grossman, 2024).

Secondly, in adults, the HRV component also reflects an alert state in response to the stimulus, but this is accompanied by a decrease in RSA, potentially associated with an increased motor response— an aspect not yet present in the neonatal period. The differing autonomic responses to music between adults and preterm infants may be ascribed to distinct levels of motor engagement and the maturity of the autonomic system. In adults, music frequently evokes motor responses, including rhythmic movement or internal motor simulation (Chen et al., 2008), here resulting in diminished HF power and RSA. This pattern indicates a transition towards sympathetic dominance, promoting active involvement and increased arousal states.

In contrast, newborns do not typically exhibit motor engagement in response to prolonged musical stimuli, showing an initial motor response to rhythm around 7 months of age (Zentner & Eerola, 2010). Instead, they tend to recruit brain areas outside the sensorimotor regions when processing instrumental sounds (Loukas et al., 2024). Consequently, although adults may demonstrate heightened arousal and motor readiness in reaction to music, preterm infants do not display similar autonomic alterations, indicative of their developmental phase.

Thirdly, the presence of a vagal cardiac control in cardiorespiratory dynamics of preterm infants and, as opposite, of a respiratory driving mechanism in adults is demonstrated. The directionality of cardiorespiratory communications remains a topic of debate, and one of the most shared hypothesis is that physiological mechanisms at the base of these interaction are bi-directional, with complex feedback loops between heart and lungs, modulated by mechanical factors, brainstem control, ANS activity, baroreceptors and chemoreceptors feedback (Dick et al., 2014; Elstad et al., 2018; Garcia III et al., 2013). Multiple independent coupling methods have been proposed and applied (Bartsch et al., 2014), RSA and Phase Coupling being just two of them. In adults, literature consistently reports a stronger influence of breath on cardiac rhythm, irrespective of the analytical approach used (Schulz et al., 2013). Both linear (Faes & Nollo, 2010; Faes et al., 2004) and nonlinear methods (Riedl et al., 2010) (Bahraminasab et al., 2008) confirm this predominant directionality (Sobiech et al., 2017) (Borovkova et al., 2022). Our findings align with these studies, as we observe a consistent directional influence from respiration to heart rate throughout the study protocol.

In newborns, however, these interactions present additional complexities, as the ANS is still rapidly developing (Lucchini et al., 2018), and RSA has been shown to correlate with developmental age and increase within the first six months (Harper et al., 1978). Studies on phase coupling indicate a developmental transition from bidirectional to unidirectional, respiration-driven interactions over the first six months of life (Mrowka et al., 2003). However, direct comparisons between studies remain challenging due to methodological differences and variations in the specific phenomena analysed. In preterm infants, these feedback mechanisms are even less developed (Chatow et al., 1995), potentially leading to distinct regulatory patterns. Understanding these interactions in premature neonates is essential for investigating the maturation of ANS-driven modulation and identifying potential disruptions in the underlying processes. Preterm infants are particularly vulnerable to cardiorespiratory complications (Raju et al., 2017), often exacerbated by the NICU environment, where external conditions significantly influence autonomic development (Patural et al., 2008; Schlatterer et al., 2022). The parasympathetic system, which plays a crucial role in cardiorespiratory interactions, undergoes significant developmental changes in the weeks following preterm birth (Mulkey & Du Plessis, 2019).

Notably, preterm infants typically exhibit a less developed or even absent RSA (Clark et al., 2012; Indic et al., 2011; Indic et al., 2008; Joshi et al., 2019; Longin, Gerstner, Schaible, Lenz, & Koenig, 2006; Rassi et al., 2005) which may have long-term health implications (Mathewson et al., 2014). The variability in RSA presence and strength reported in the literature (Clark et al., 2012; Joshi et al., 2019; Longin, Gerstner, Schaible, Lenz, & König, 2006) may stem from its transient nature, suggesting that RSA is an evolving mechanism rather than a fully established one at preterm birth.

Additionally, studies on preterm infants in the NICU have reported bidirectional interactions at both nonlinear (Porta-García et al., 2023) and linear levels (Indic et al., 2011), supporting the notion that cardiorespiratory coordination in neonates differs from the respiration-driven interactions observed in adults. Our findings further reinforce this hypothesis, demonstrating that very preterm infants do not yet exhibit the stable respiratory-led interaction pattern characteristic of mature individuals. This aligns with findings by Joshi et al. (Joshi et al., 2019), who reported a similar heart-driven interaction pattern using comparable methodologies. Furthermore, our results highlight significant differences between premature infants and adults, with preterm infants displaying stronger heart-to-respiration coupling compared to the mature respiration-to-heart influence seen in adults.

Finally, some studies suggest that intervention protocols could positively influence the development of these mechanisms (Bloch-Salisbury et al., 2009; Indic et al., 2011). The present study supports this perspective, as we observed an increase in RSA following musical exposure in preterm infants, along with a shift in dynamics during music stimulation from a cardiac-led interaction to a more balanced coupling strength.

### Limitation

To assess the progressive maturation of the cardiorespiratory system through this multi-level analysis of responses to musical stimuli, it is essential to cover critical ages across development. First, it is fundamental to include a term reference population in the coming research to disentangle the specific effects of prematurity. Furthermore, two distinct age ranges in preterm development—before 32 weeks of GA and term age—will be particularly interesting to be investigated. The progressive maturation of the cardiorespiratory system during a preterm infant’s stay in the NICU is influenced by both intrinsic maturation mechanisms and environmental factors. Determining whether a music intervention could progressively support this maturation is crucial for developing early intervention guidelines in this at-risk population.

## CONCLUSION

Our findings offer new insights into the development of cardiorespiratory interactions in preterm infants, highlighting the similarities and differences of their autonomic regulation compared to adults. While HRV responses to music were similar across both groups, a more detailed analysis of RSA and phase coupling revealed fundamental differences in the underlying mechanisms. These results carry interesting implications for neonatal care, particularly in the integration of music-based interventions within the NICU environment. By deepening our understanding of how music influences autonomic regulation in preterm infants, this study reinforces the potential benefits of auditory enrichment in neonatal care. Future work should also assess the practical implementation of tailored soundscapes in the NICU to promote optimal autonomic and neurophysiological development, potentially reducing the risks associated with autonomic dysfunction in this vulnerable population.

## Acknowledgements

This work was supported by the Swiss National Science Foundation (FNS)) (Principal Investigator: Prof. Petra Hüppi, Grant ID: F02-12516), as well as the Dora Foundation, Prim’Enfance Foundation, K Foundation, and the Von Meissner Foundation

## STAR Methods

| REAGENT or RESOURCE | SOURCE | IDENTIFIER |
| --- | --- | --- |
| Biological samples |  |  |
| 22 infants | Neonatal Intensive Care Unit (NICU) at the University Hospitals of Geneva, Switzerland |  |
| 20 adults | University of Geneva |  |
| Software and algorithms |  |  |
| MATLAB (R2021b) | MathWorks Inc, Natick, MA, USA | https://it.mathworks.com/products/matlab |
| Kubios HRV Standard (v2.2) | (Tarvainen et al., 2014) | https://www.kubios.com/ |

## RESOURCE AVAILABILITY

### Materials availability

This study did not generate new unique reagents.

### Data and code availability

This paper does not report original code.

## EXPERIMENTAL MODEL AND STUDY PARTICIPANT DETAILS

### Ethics statement

The Swiss Research Ethics Committee approved the study (BASEC no. 2023-00718), and written informed consent was obtained from all parents

### Human participants

A total of 22 infants born before 32 weeks of gestation were recruited from the Neonatal Intensive Care Unit (NICU) at the University Hospitals of Geneva, Switzerland. One infant was excluded due to excessive fussiness, and two were excluded due to instrumentation issues. Infant’s demographic data for the final population are presented in Table 6.

**Table 6.** Characteristics of the infant population, Note. GA = gestational age.

| Characteristics | Value |
| --- | --- |
| Population, N | 18 |
| GA at birth (weeks), (mean ± std) | 27,63 ± 2.24 |
| Birthweight, (g), (mean ± std) | 974,56 ± 379,66 |
| GA at test (week), (mean ± std) | 37.73 ± 0.80 |

Additionally, a cohort of 20 adults was recruited to undergo the same music protocol as the infants. One adult was excluded due to instrumentation issues. Adult’s demographic data are reported in Table 7.

**Table 7.**
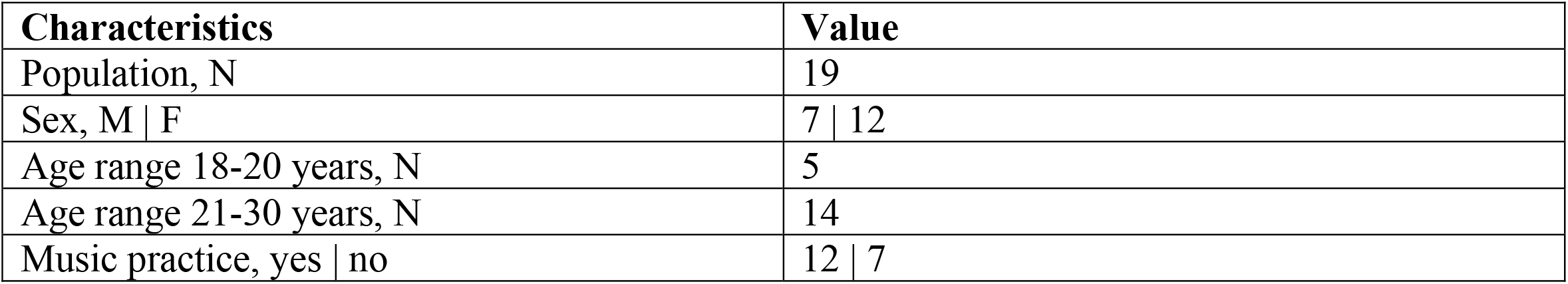
Characteristics of the adult population.

| Characteristics | Value |
| --- | --- |
| Population, N | 19 |
| Sex, M \| F | 7 \| 12 |
| Age range 18-20 years, N | 5 |
| Age range 21-30 years, N | 14 |
| Music practice, yes \| no | 12 \| 7 |

## METHOD DETAILS

### EXPERIMENTAL PROCEDURE

While in intensive care, infants were exposed to music intervention via headphones Sennheiser PXC 250-II (Sennheiser electronic GmbH & Co.KG, Wedemark, Germany), adapted to the size of the baby’s head. During the whole protocol physiological signals were acquired. A representation of the set-up can be seen in Figure 10.

**Figure 9.**
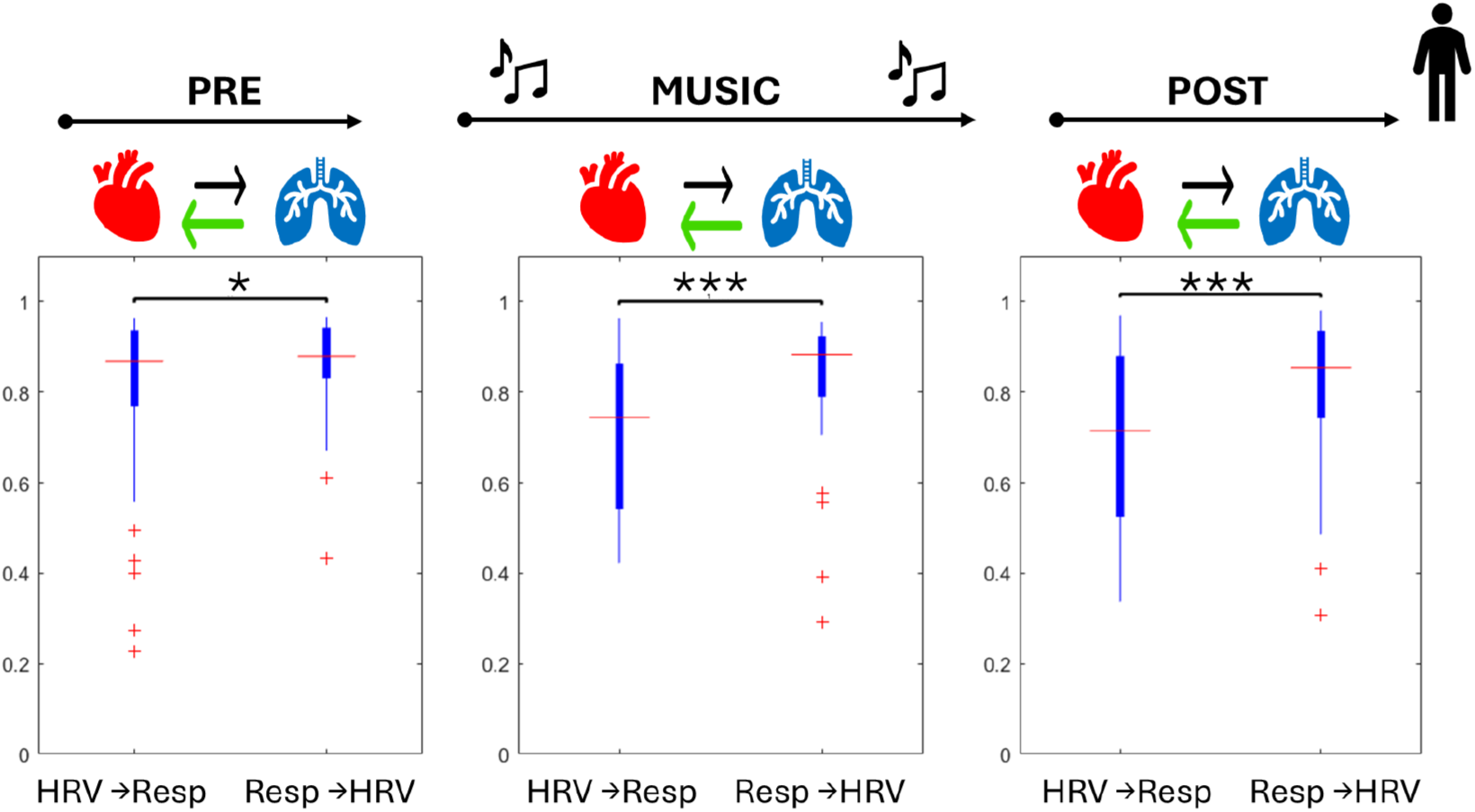
Directionality analysis for adults during the protocol. Boxplots depict the aggregation of cross correlation BPRSA values for each direction. Significance thresholds are represented by “* = p <0.05”, “** = p < 0.01”. In the top line protocol phases are represented, and arrow depicting a significantly stronger directionality is highlighted in yellow.

**Figure 10.**
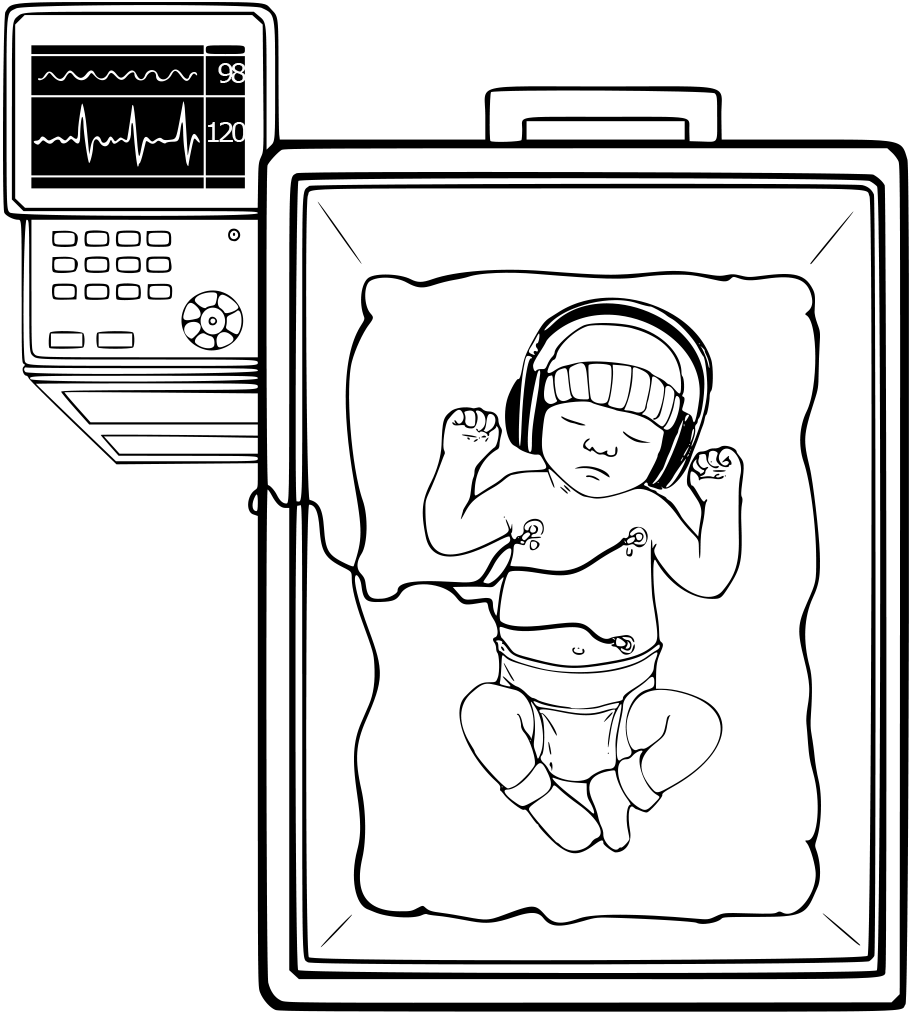
Representation of the acquisition set-up. The infant was placed in the NICU incubator, wearing size-adjusted headphones, while physiological signals were acquired using electrodes in a 3-lead configuration.

**Figure 11.**
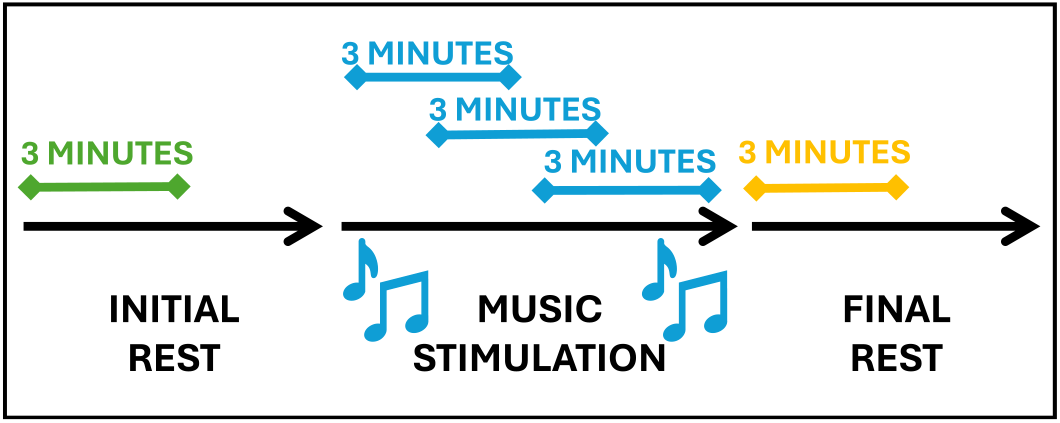
Temporal representation of the protocol steps

A baseline period of at least 3 minutes was acquired before and after the music stimulation, defined as a period in which infant was wearing the headphones without music was otherwise unstimulated. The music stimulation consisted in 6 minutes of stereophonic musical excerpts in B major, specifically created by the musician Andreas Vollenweider, with a sound level from 30dBA for the background to 65dBA for the peak with bells.

The music has been used in similar protocol in (Lordier, Meskaldji, et al., 2019), (Barcos-Munoz et al., 2024). The music piece is built on a harmonic foundation of human voices, occasionally enriched by harp, punji, and bells. The landscape is stereophonic, with instrument appearing intermittently. The main elements are the harp, introducing a soft texture and melodic emphasis, while the punji, a traditional Indian flute, carrying the principal melody and bells providing sporadic percussive accents in the harmonic structure.

### Electrophysiological recording

Infant physiological signals were recorded using standard cardiorespiratory monitoring with the IntelliVue MP40 monitor (Philips Medical System, Eindhoven, Netherlands). The ECG signal was then automatically recorded using the IxTrend 2.1 software (Ixellence GmbH, Wildau, Germany; ECG was acquired at 500 Hz while the respiratory signal was acquired at 62.5 Hz. The adult’s ECG signals were recorded using the ANT Neuro waveguard™ system at a sampling frequency of 1000 Hz.

## DATA PREPROCESSING

### ECG pre-processing

The interbeat interval series were derived from the ECG signals using the Kubios 2.2 software (Tarvainen et al., 2014), following visually inspection and artifact correction. The resulting series were detrended with the smoothness priors’ approach (Tarvainen et al., 2002), setting λ = 500, and resampled at 4Hz using a piecewise cubic spline interpolation. The resulting uniformly sampled series will henceforth be called HRV signals.

### Infants’ respiratory signals preprocessing

Infants’ respiratory signals were pre-processed following (Zhang et al., 2021). Signals were downsampled at 16 Hz filtered with a 5th-order Butterworth filter ([0.05–2] Hz), baseline-corrected using a 1.2s median trend removal, resampled to 4 Hz, and normalized via z-score.

### Adults’ respiratory signals preprocessing

For adults, since breath signals were not available, the respiratory activity was derived from the ECG using the slope-range method (Kontaxis et al., 2020). The ECG-derived respiratory signal (ECG-DR) served as a validated surrogate for the respiratory signal in adults (Varon et al., 2019). ECG signals were filtered with a twelve-order high-pass Butterworth filter (0.8 Hz) to remove baseline wander and a twelve-order low-pass Butterworth filter (30 Hz) to eliminate high-frequency noise. The slope range was defined as the difference between the maximum up-slope and the minimum down-slope within the QRS interval and used to represent respiratory activity, for details see (Kontaxis et al., 2020). An outlier correction procedure was performed according to (Bailon et al., 2006), and signals were resampled at 4 Hz using cubic spline interpolation to match the HRV series.

## DATA ANALYSIS

### Window selection

Autonomic activity and cardiorespiratory coupling were analyzed in three stages: before, during, and after musical elicitation. Three-minute windows were used: initial baseline, three overlapping windows during stimulation, and post condition (see Figure 10). For music elicitation, the average of the feature values over the three overlapping windows was computed to characterize the global effect.

### HRV frequency analysis

The autonomic modulation on HRV was investigated by using the standard frequency analysis and adding the information obtained from the application of the Orthogonal Subspace Decomposition (OSP) (Varon et al., 2019; Widjaja et al., 2013).

#### Standard frequency analysis

The power density spectrum (PDS) of HRV signals was computed for each three-minute window using Welch’s method. HRV signal power was computed in two frequency bands: LF and HF, with age-specific ranges—[0.04–0.15] Hz (LF) and [0.15–0.4] Hz (HF) for adults, and [0.02–0.2] Hz (LF), and [0.2–1.5] Hz (HF) for infants, according to (Filippa et al., 2022; Rosenstock et al., 1999). LF and HF power were calculated, along with the LF/HF ratio, which serves as an index of physiological arousal, while HF power reflects parasympathetic modulation (“Heart rate variability,” 1996).

#### Orthogonal Subspace Decomposition (OSP)

To identify factors influencing HRV changes, the OSP method (Varon et al., 2019; Widjaja et al., 2013) was applied. OSP separates respiratory components linearly related to HRV from residual ones, such as sympathetic and vagal modulations unrelated or non-linearly related to respiration. This technique produces two signals: one influenced by the respiratory activity and another for residual activity.

The HRV signal is projected into two orthogonal subspaces, representing the respiratory activity and the residual activity respectively, as follows. Given a respiratory signal X = x(1), …, x(N), and the HRV signal Y = y(1), …, y(N), and the model order m, a subspace V defined by all variations in X is constructed using X and its delayed version from 1 to m samples:

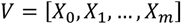

The vectors X_d_ are defined as:

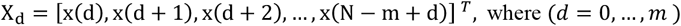

Using this subspace, a projection matrix P is defined as:

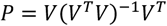

Then Y is projected over the respiratory subspace using P, and Y_X_, the HRV component linearly related to respiration is obtained as:

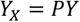

The residual is calculated as the orthogonal component:

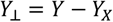

The model order m has been defined as the minimum number of delays obtained considering the minimum description length (MDL) principle and the Akaike Information Criterion (AIC).

Once the two components were defined, the power in the previously described frequency bands was calculated for both components in order to characterize the respective phenomena.

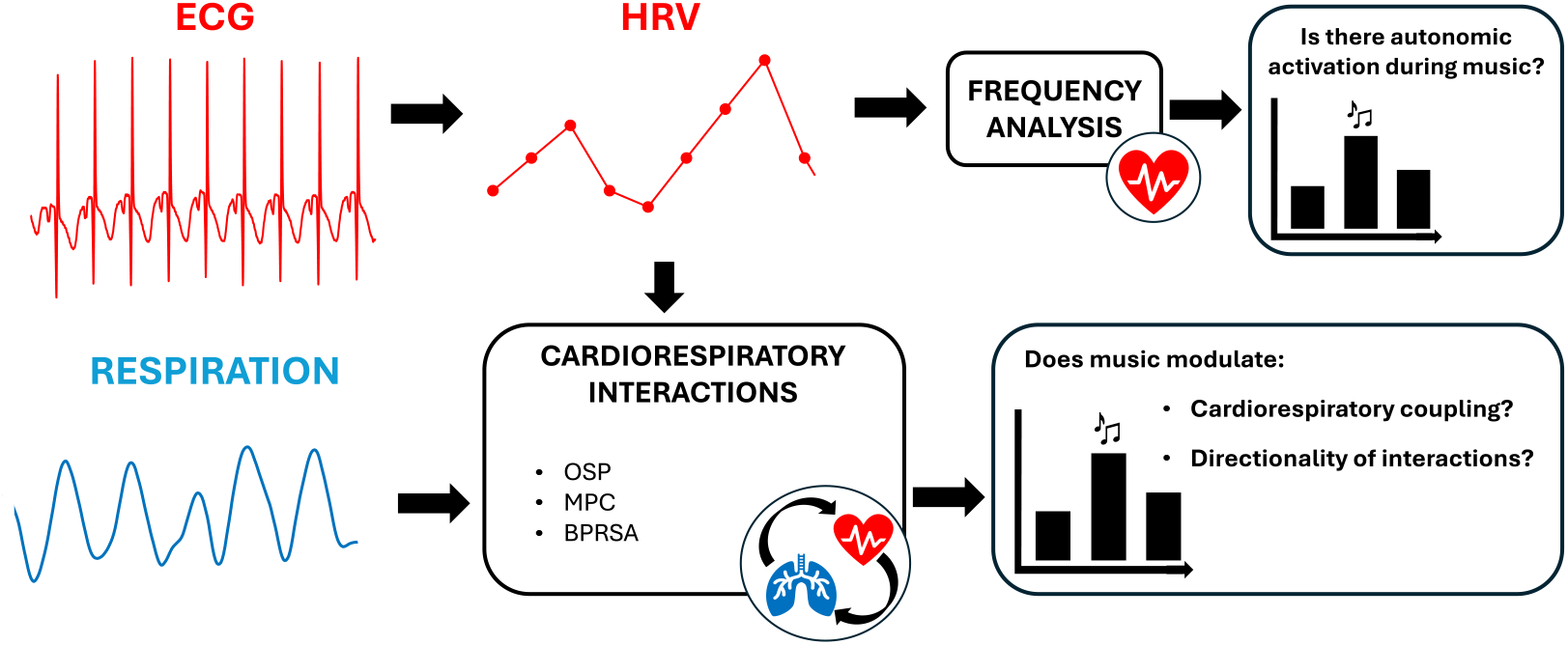

#### Cardiorespiratory analysis

Combining the information extracted from the HRV and the respiratory signals, we applied three approaches to investigate the cardiorespiratory interaction: the relative power of the respiratory component obtained through the OSP (Varon et al., 2019; Widjaja et al., 2013), the Mean Phase Coherence (MPC) (Lanata et al., 2016), and the Bivariate Phase-Rectified Signal Averaging (BPRSA) (Schumann et al., 2008).

#### OSP-derived relative power of the respiratory component

The percentage of the power related to the respiration component with respect to the total power of the HRV signal (hereinafter P_x_) was computed as an index of strength of cardiorespiratory coupling, quantifying the amount of information shared between respiration and heart rate variability dynamics (Morales et al., 2021; Varon et al., 2019).

The metric P_x_ is calculated as:

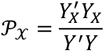

This approach has been reported as one of the most effective to quantify RSA strength (Morales et al., 2021) and used to assess the phenomena in clinical evaluations (Armañac-Julián et al., 2021).

#### Mean Phase Coherence (MPC)

The Mean Phase Coherence (MPC) quantifies the phase synchronization of two time-series regardless of their amplitude.

This synchronisation has been observed in chaotic systems and coupled complex oscillators (Parlitz et al., 1996; Rosenblum et al., 1996). According to the previous literature, the MPC is suitable for application to time series expressing coupled physiological dynamics, such the heart and respiration activities, as they display clear nonlinear, complex behaviour (Goldberger et al., 2002) and can act as weakly coupled complex systems (Schäfer et al., 1998).

The MPC metric has been computed as described in (Lanata et al., 2016). Given two time-series x and y:

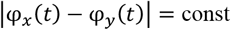

with *φ*_*x*_ (*t*) and *φ*_*y*_ (*t*) representing the instantaneous phases of x and y respectively, the final metrics is computed as follows

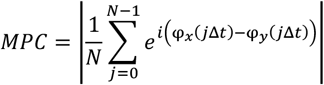

where t is the time resolution, N is the sample number of each signal.

Phase-synchronization of cardiorespiratory series has been previously observed in the literature in adults (Schäfer et al., 1998), and in newborn infants (Mrowka et al., 2000).

#### Bivariate Phase-Rectified Signal Averaging (BPRSA)

The Bivariate Phase-Rectified Signal Averaging approach (Schumann et al., 2008) quantifies the inter-relation between two signals, independently from non-stationarity and noise.

It detects anchor points x_*i*1_, x_*i*2_, …, x_*i*M_ on a trigger signal X and their respective timing and averages corresponding fixed-length windows (2L) on target signal Y, yielding a modulated signal. The result is a signal of length 2L, representing the modulations on the target signal attributed to periodicities in the trigger. Given a total of M anchor points, the trend is defined as:

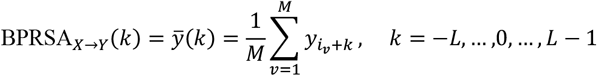

Respiratory and HRV time series (without detrending) were analyzed using L=55 samples in both interaction directions.

For the first direction For HRV as the trigger, anchor points corresponded to heart rate acceleration and deceleration, where deceleration was defined as:

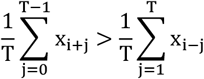

given x the value of the HRV serie, as such a deceleration in cardiac activity corresponds to an increase in the RR time intervals. For these computations T has been set to 1, as done in (Joshi et al., 2019).

For the reverse direction, the respiration time series has been used as a trigger signal, selecting two types of anchor points, i.e., the maximum and the minimum of each respiratory cycle.

Then, the strength of the coupling between the trigger and the target has been computed using a novel approach, according to (Montero-Nava et al., 2020) (Esquivel-Arizmendi et al., 2020), using both linear (maximum absolute value of normalized cross-correlation) and nonlinear (Mutual Information, MI) methods (Fraser & Swinney, 1986) via the Kernel density estimation method (Moon et al., 1995).

#### Statistical analysis

Statistical analysis was used to compare the markers of autonomic dynamics during the elicitation protocol phases. Normality was checked using the Lilliefors test. Due to non-Gaussian distributions, the Friedman test compared changes across phases (initial rest, music elicitation, final rest). When statistically significant differences were found (Friedman’s p-value < 0.05), post-hoc pairwise Wilcoxon tests with Bonferroni correction were computed.

### Directionality assessment

For the BRSA analysis, differences in coupling directionality were evaluated by comparing the coupling strength between HRV → respiration during acceleration/deceleration and respiration → HRV during minimum/maximum impedance for each phase. In the absence of a specific hypothesis for selecting the triggers, acceleration/deceleration for HRV → respiration and maximum/minimum for respiration → HRV were aggregated. This comparison aimed to explore directional dynamics (not causality) using both cross-correlation and Mutual Information (MI). Wilcoxon signed rank tests for paired samples were used, with Bonferroni correction applied for multiple comparisons.

---

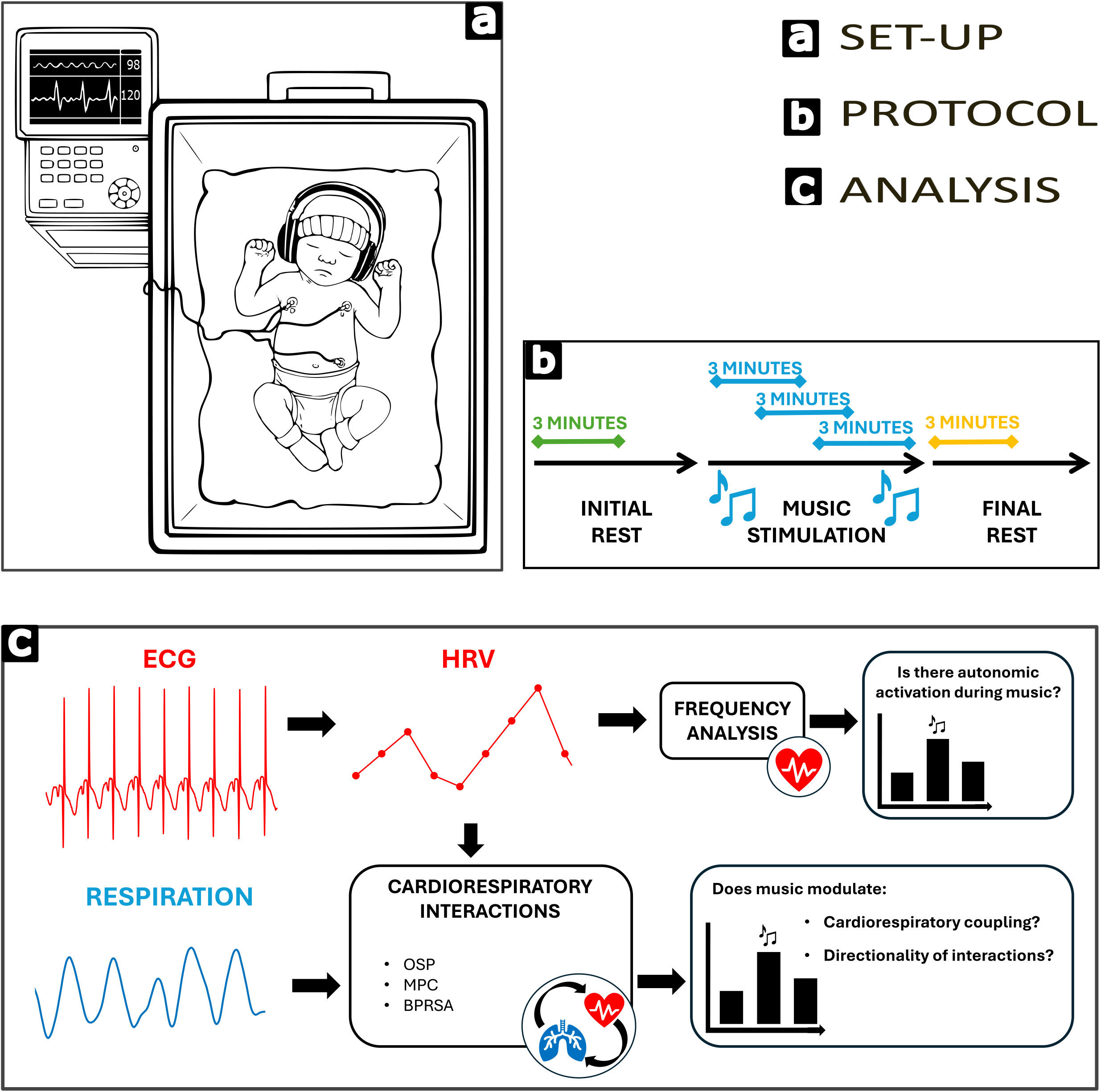

